# Limiting the priming dose of a SARS CoV-2 vaccine improves virus-specific immunity

**DOI:** 10.1101/2021.03.31.437931

**Authors:** Sarah Sanchez, Nicole Palacio, Tanushree Dangi, Thomas Ciucci, Pablo Penaloza-MacMaster

**Affiliations:** Department of Microbiology-Immunology, Feinberg School of Medicine, Northwestern University, Chicago, IL 60611, USA.; David H. Smith Center for Vaccine Biology and Immunology, University of Rochester, Rochester, NY 14642, USA.; Department of Microbiology and Immunology, Center for Vaccine Biology and Immunology, University of Rochester, Rochester, NY 14642, USA.

**Author notes:** **Correspondence:** Pablo Penaloza-MacMaster.

## Abstract

Since late 2019, SARS-CoV-2 has caused a global pandemic that has infected 128 million people worldwide. Although several vaccine candidates have received emergency use authorization (EUA), there are still a limited number of vaccine doses available. To increase the number of vaccinated individuals, there are ongoing discussions about administering partial vaccine doses, but there is still a paucity of data on how vaccine fractionation affects vaccine-elicited immunity. We performed studies in mice to understand how the priming dose of a SARS CoV-2 vaccine affects long-term immunity to SARS CoV-2. We first primed C57BL/6 mice with an adenovirus-based vaccine encoding SARS CoV-2 spike protein (Ad5-SARS-2 spike), similar to that used in the CanSino and Sputnik V vaccines. This prime was administered either at a low dose (LD) of 10^6^ PFU or at a standard dose (SD) of 10^9^ PFU, followed by a SD boost in all mice four weeks later. As expected, the LD prime induced lower immune responses relative to the SD prime. However, the LD prime elicited immune responses that were qualitatively superior, and upon boosting, mice that were initially primed with a LD exhibited significantly more potent immune responses. Overall, these data demonstrate that limiting the priming dose of a SARS CoV-2 vaccine may confer unexpected benefits. These findings may be useful for improving vaccine availability and for rational vaccine design.

## Introduction

Several vaccines are currently being used in humans to prevent COVID-19. Among these, adenovirus-based vaccines have shown potent protection against severe COVID-19. These vaccines are based on adenovirus serotype 5, adenovirus serotype 26, and chimpanzee adenovirus (ChAdOx1). However, there is not a sufficient number of vaccine doses available to immunize the entire world population, which has motivated discussions about administering partial vaccine doses to increase the number of people who receive the vaccine. Nevertheless, there is little information on how vaccine fractionation affects long-term immunity to SARS CoV-2.

Phase I vaccine trials typically involve dose-escalation studies comparing a range of vaccine doses in groups of people who receive the same dose of the vaccine during the prime and the boost. However, they do not typically evaluate “intra-group dose escalation,” in which individuals would first receive a prime with a low dose, and then a boost with a higher dose. We performed studies in mice to determine the immunological effect of intra-group vaccine dose escalation. Our data show that limiting the priming dose of a SARS CoV-2 vaccine may confer an unexpected qualitative benefit to T cell responses and antibody responses.

## Results

### Low dose (LD) vaccine prime elicits T cells with high anamnestic capacity

We first primed C57BL/6 mice intramuscularly with an adenovirus vector expressing SARS CoV-2 spike protein (Ad5-SARS-2 spike), either at a low dose **(LD) (10^6^ PFU)** or a standard dose **(SD) (10^9^ PFU)**. After four weeks, mice were boosted with the standard dose and CD8 T cell responses were analyzed by MHC tetramer binding assays (Fig. 1A). We tracked a CD8 T cell response against an epitope (VNFNFNGL) that is highly conserved among bat and SARS-like coronaviruses, including SARS CoV-1, SARS CoV-2, RatG13, HKU3, WIV1, WIV16, RsSHC014, Rs3367, Shaanxi2011, Rm1/2004, YN2018B, SC2018, HuB2013, GX2013, and BM48-31/BGR/2008/Yunnan2011, among other coronaviruses. We will refer to this conserved CD8 T cell response as SARS CoV-2 specific response, or K^b^ VL8 for simplicity purposes, where <u>V</u> represents the first amino acid, <u>L</u> represents the last amino acid, and <u>8</u> represents the number of amino acids in the sequence.

**Figure 1.**
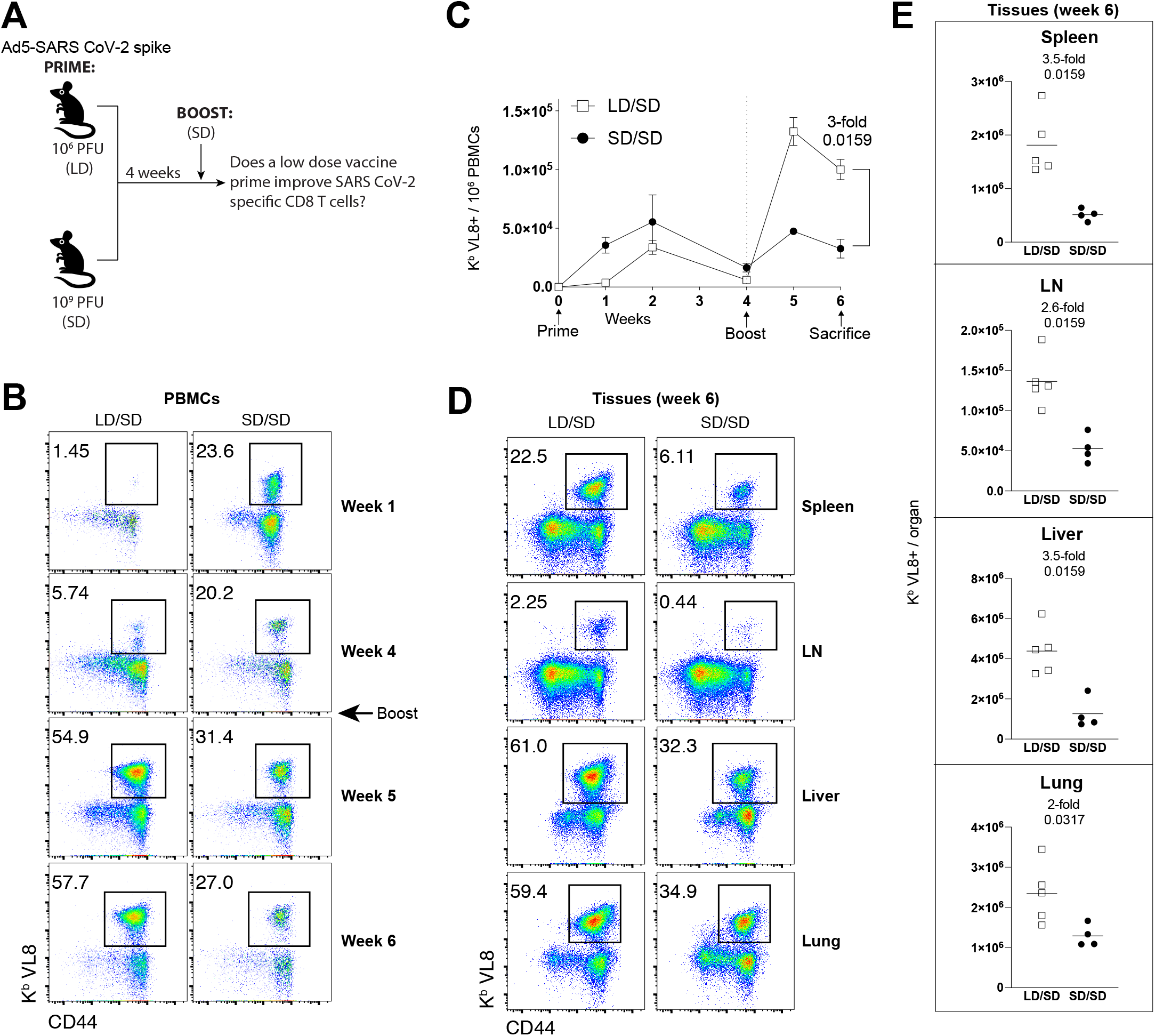
A LD/SD vaccine regimen elicits superior CD8 T cells compared to a SD/SD vaccine regimen. **(A)** Experimental approach for evaluating how the priming dose of an Ad5-SARS-2 spike vaccine affects CD8 T cell responses in C57BL/6 mice. **(B)** Representative FACS plots showing the frequencies of SARS CoV-2-specific CD8 T cells (K^b^ VL8+) in PBMCs. **(C)** Summary of SARS CoV-2-specific CD8 T cell responses in PBMCs. **(D)** Representative FACS plots showing the frequencies of SARS CoV-2-specific CD8 T cells (K^b^ VL8+) in tissues. **(E)** Summary of SARS CoV-2-specific CD8 T cell responses in tissues. Data are from one experiment with n=5 per group. Experiment was repeated two additional times with similar results. Indicated *P* values were determined by Mann-Whitney U test. Error bars represent SEM.

Initially, priming with a LD resulted in lower SARS CoV-2 specific CD8 T cell responses, relative to priming with a SD (Fig. 1B). This result is consistent with the notion that the level of adaptive immune responses following adenovirus vaccination is dosedependent (*1*). However, an unexpected effect was observed after the booster immunization four weeks after. Mice that were initially primed with a LD exhibited significantly greater CD8 T cell expansion upon boosting, relative to mice that were initially primed with the higher SD (Fig. 1B-1C). More robust CD8 T cell recall expansion in the LD/SD regimen was also observed in tissues (Fig. 1D-1E).

SARS CoV-2-specific CD8 T cells induced by the LD/SD regimen exhibited enhanced CD107a degranulation and IFNγ expression (Fig. 2A-2C). Moreover, there was a pattern of improved CD4 T cell responses in the LD/SD regimen relative to the SD/SD regimen, but the difference was not statistically significant (Fig. 2D). SARS CoV-2-specific CD8 T cells induced by the LD/SD regimen showed more robust granzyme B and Ki67 expression relative to the SD/SD regimen (Fig. 2E-2G), suggesting enhanced cytotoxicity and proliferation upon boosting. Collectively, these results demonstrate that limiting the priming dose elicits SARS CoV-2-specific T cell responses with superior anamnestic and functional capacity.

**Figure 2.**
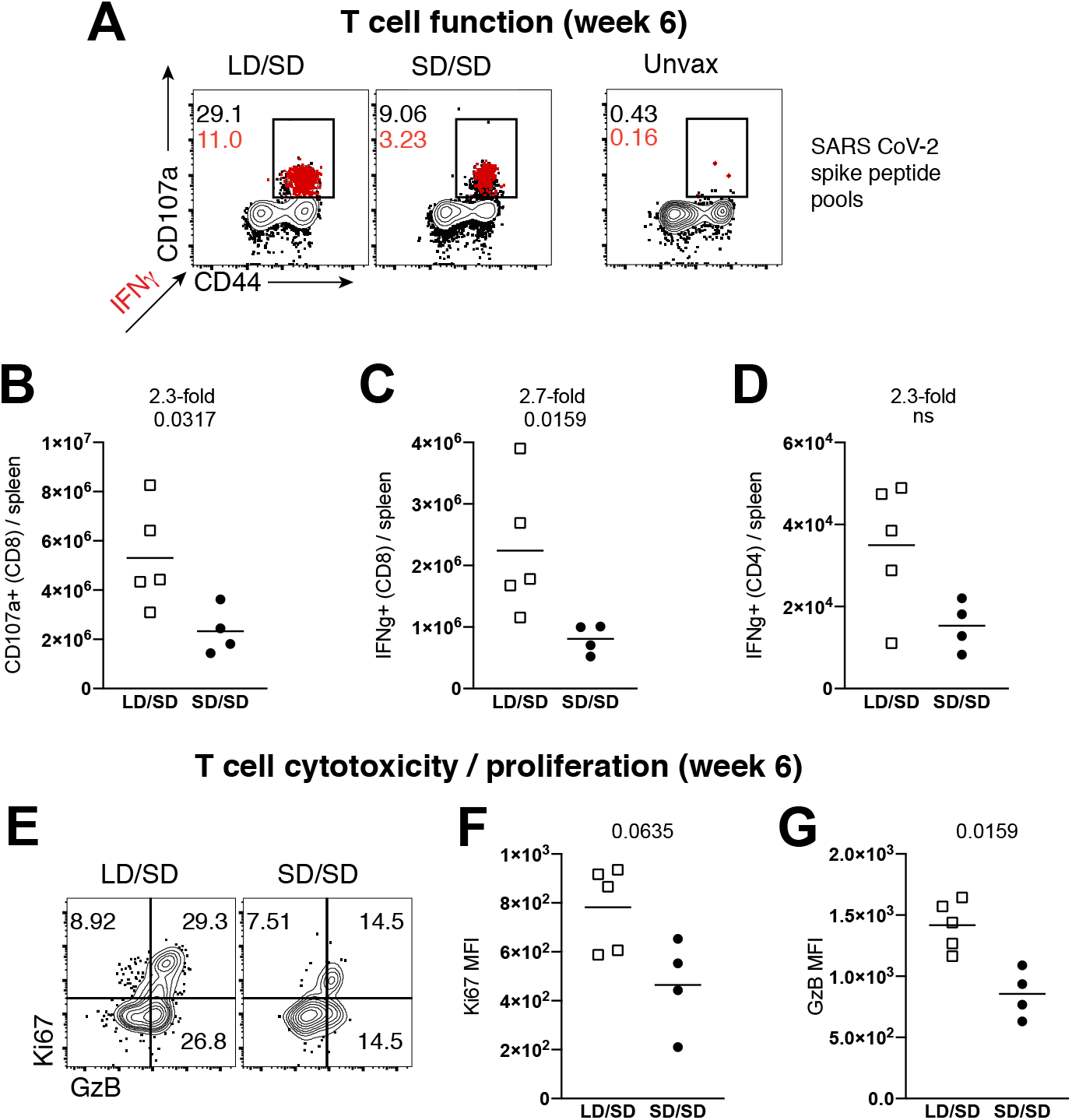
A LD/SD vaccine regimen elicits more functional CD8 T cell responses compared to a SD/SD vaccine regimen. In panels A-D, splenocytes were incubated with overlapping SARS CoV-2 peptide pools for 5 hr at 37°C in the presence of GolgiStop and GolgiPlug. **(A)** Representative FACS plots showing the frequencies of cytokine expressing SARS CoV-2-specific CD8 T cells. **(B)** Summary of SARS CoV-2-specific CD8 T cells that express the degranulation marker CD107a. **(C)** Summary of SARS CoV-2-specific CD8 T cells that express IFNγ. **(D)** Summary of SARS CoV-2-specific CD4 T cells that express IFNγ. **(E)** Representative FACS plots showing the frequencies of granzyme B and Ki67 expressing CD8 T cells. **(F)** Summary of Ki67 expression. **(G)** Summary of granzyme B expression. Panels E-G are gated on K^b^ VL8+ cells (SARS CoV-2-specific). Data from panels F-G are indicated as mean fluorescence intensity (MFI). All data are from spleen. Data are from one experiment with n=4-5 per group. Experiment was repeated once with similar results. Indicated *P* values were determined by Mann-Whitney U test.

### Effects of vaccine dose on T cell differentiation

The data above suggested qualitative differences in T cell responses following a single shot with either a low or a standard dose of vaccine. It is known that after an initial antigen encounter, T cells differentiate into distinct subsets, including short-lived effector cells (Teff), effector memory cells (Tem), and central memory cells (Tcm). Teff and Tem subsets exhibit rapid cytotoxicity, but are short-lived. On the other hand, the Tcm subset is long-lived and exhibits enhanced recall capacity (*2*). To evaluate if the priming dose affected the differentiation of these T cells subsets, we FACS-sorted SARS CoV-2-specific CD8 T cells at week 4 post-vaccination (prime only), and performed single-cell RNA-sequencing (scRNA-seq). Interestingly, SARS CoV-2 specific CD8 T cells elicited by a LD and a SD clustered differently, suggesting unique transcriptional signatures (Fig. 3A). CD8 T cells after a LD prime showed lower transcription of genes associated with effector function, such as *Prdm1, Tbx21, Id2, Ifng, Gzmb, Gzma, Prf1,* and genes associated with terminal differentiation, such as *Klrg1* and *Zeb2* (Fig. 3B). Overall, the LD was associated with lower expression of genes associated with terminal effector differentiation (Fig. 3C)(*3*). Importantly, CD8 T cells elicited by a LD showed higher transcription of genes associated with long-lived memory T cells, such as *Tcf7, Id3, Bcl2, Ccr7, Sell (CD62L)* and *Il7r* (CD127) (Fig. 3B). Taken together, although the LD prime elicited a lower number of CD8 T cells, most of those CD8 T cells exhibited a Tcm signature with low terminal effector differentiation.

**Figure 3.**
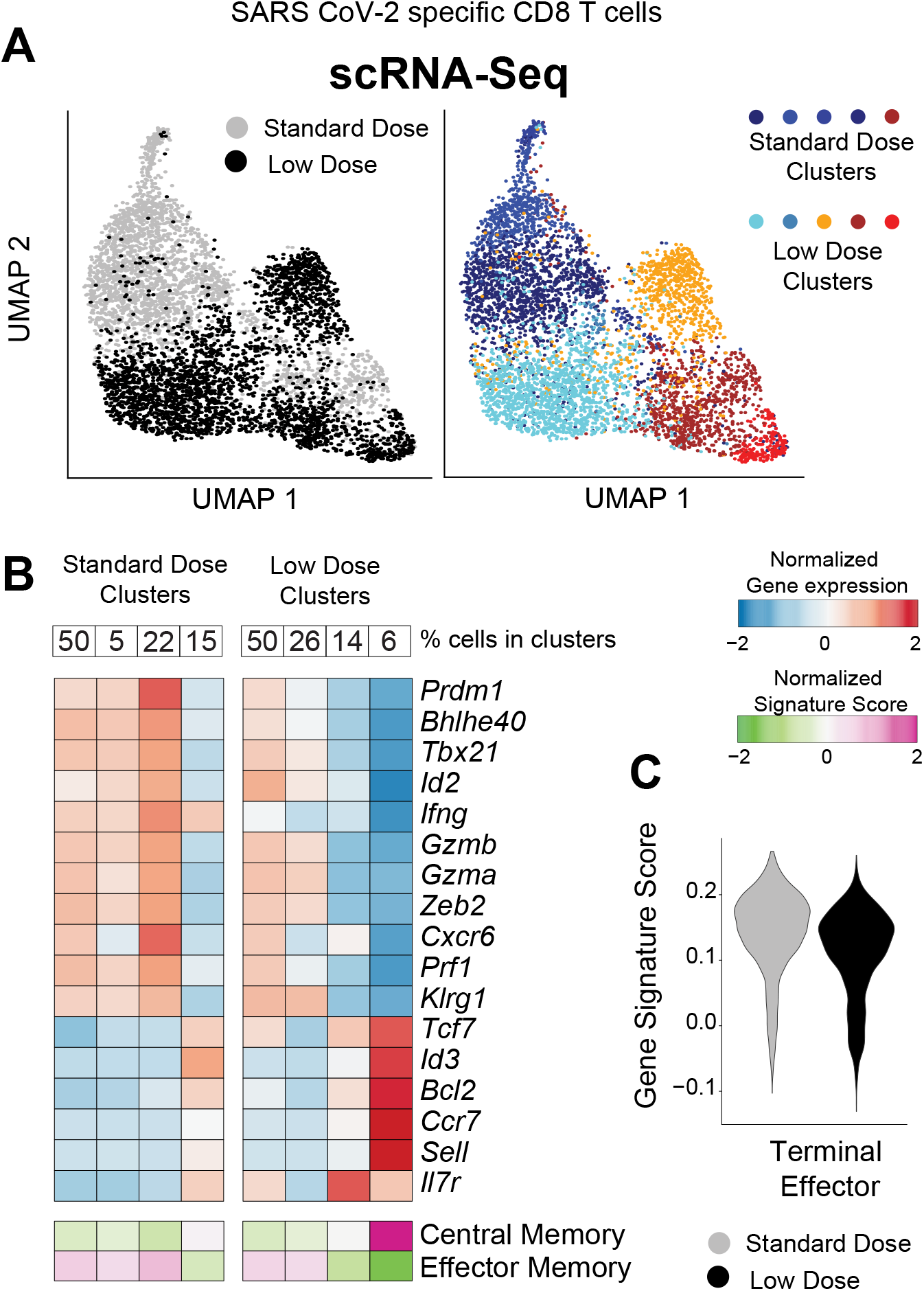
Single cell RNA-seq analyses demonstrate that a low dose prime favors central memory CD8 T cell differentiation. Mice were immunized with 10^6^ or 10^9^ PFU of Ad5-SARS-2 spike, and at day 28, splenic CD8 T cells were MACS-sorted. Subsequently, live, CD8+, CD44+, K^b^ VL8+ cells were FACS-sorted to ~99% purity for scRNA-seq. **(A)** UMAP plots showing populations colored by regimen (left plot). Standard and low dose cells were cluster separately and UMAP (right plot) shows unsupervised cell clusters. **(B)** Heatmap showing row-standardized expression of selected effector and memory genes (middle rows) or gene signatures (bottom rows). For each population, percentages of cells in each cluster are indicated (top row). **(C)** Violin plot showing the normalized expression of the Terminal Effector signature in the Standard and Low dose populations.

These data suggested that limiting the priming dose has a profound effect on T cell differentiation, and we then interrogated whether these effects on T cell differentiation could be due to distinct TCR usage. To answer this question, we performed single-cell T cell receptor (TCR)-seq on SARS CoV-2 specific CD8 T cells at week 4 post-prime, and then we compared TCR clonotypes in the two groups. In both LD and SD, TCR usage and diversity were similar and characterized by an oligoclonal expansion and a major bias for Vβ11 usage (encoded by *TRBV16* gene) (62.15% or 80.68%, respectively) (Fig. 4A-4B). Of note, we observed the presence of a dominant public clone TCRα7 TCRβ11 (*TRAV7-5/TRBV16*), sharing identical CDR3 regions (CAVIASSSFSKLVF, CASSLLGGRDTQYF) in both groups. Based on this dominant TCR sequence, we are currently creating a TCR transgenic mouse expressing the cross-reactive VL8 TCR (TCRα7 TCRβ11), which will allow one to more rigorously study CD8 T cell responses across multiple sarbecoviruses in mice. Overall, our scTCR-seq data suggested that vaccine dose did not significantly alter TCR clonotypes and viral epitope recognition at the TCR level.

**Figure 4.**
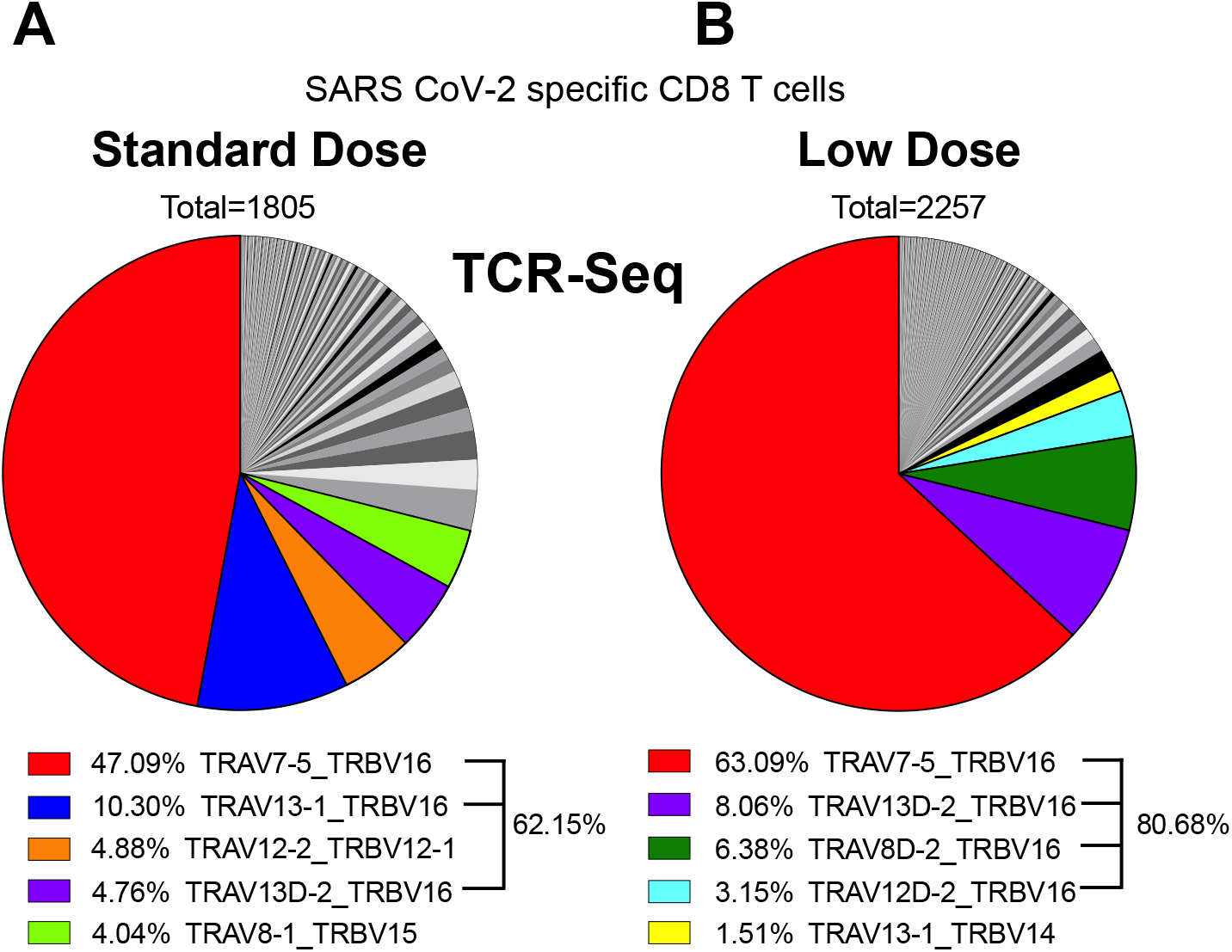
Single cell TCR-seq analyses demonstrate that the prime dose does not alter public TCR clonotypes. Mice were primed with a LD (10^6^ PFU) or a SD (10^9^ PFU) of Ad5-SARS-2 spike, and at day 28, splenic CD8 T cells were MACS-sorted. Subsequently, live, CD8+, CD44+, K^b^ VL8+ cells were FACS-sorted to ~99% purity for scTCR-Seq. Pie chart showing the distribution of TCRa and TCRb gene usage after SD prime **(A)** and LD prime **(B).** Total number above the pie chart show the number of single cells selected for the analyses, and different colors highlight the top 5 TCR usages and their relative proportion in each population.

Next, we immunophenotyped SARS CoV-2-specific CD8 T cells at week 4 post-prime to confirm the gene expression profile results. Consistent with the gene expression profiling, CD8 T cells generated by a LD prime exhibited more pronounced central memory markers, such as CD62L and CD127 (Fig. 5A-5D). In addition, CD8 T cells induced by a LD prime showed higher levels of the CD44 activation marker (Fig. 5E) and lower levels of the inhibitory PD-1 receptor (Fig. 5F), relative to CD8 T cells induced after a SD prime. Taken together, our functional data, transcriptomics data, and phenotypic data show that an initial antigen encounter has profound long-term effects on T cell differentiation following SARS CoV-2 vaccination.

**Figure 5.**
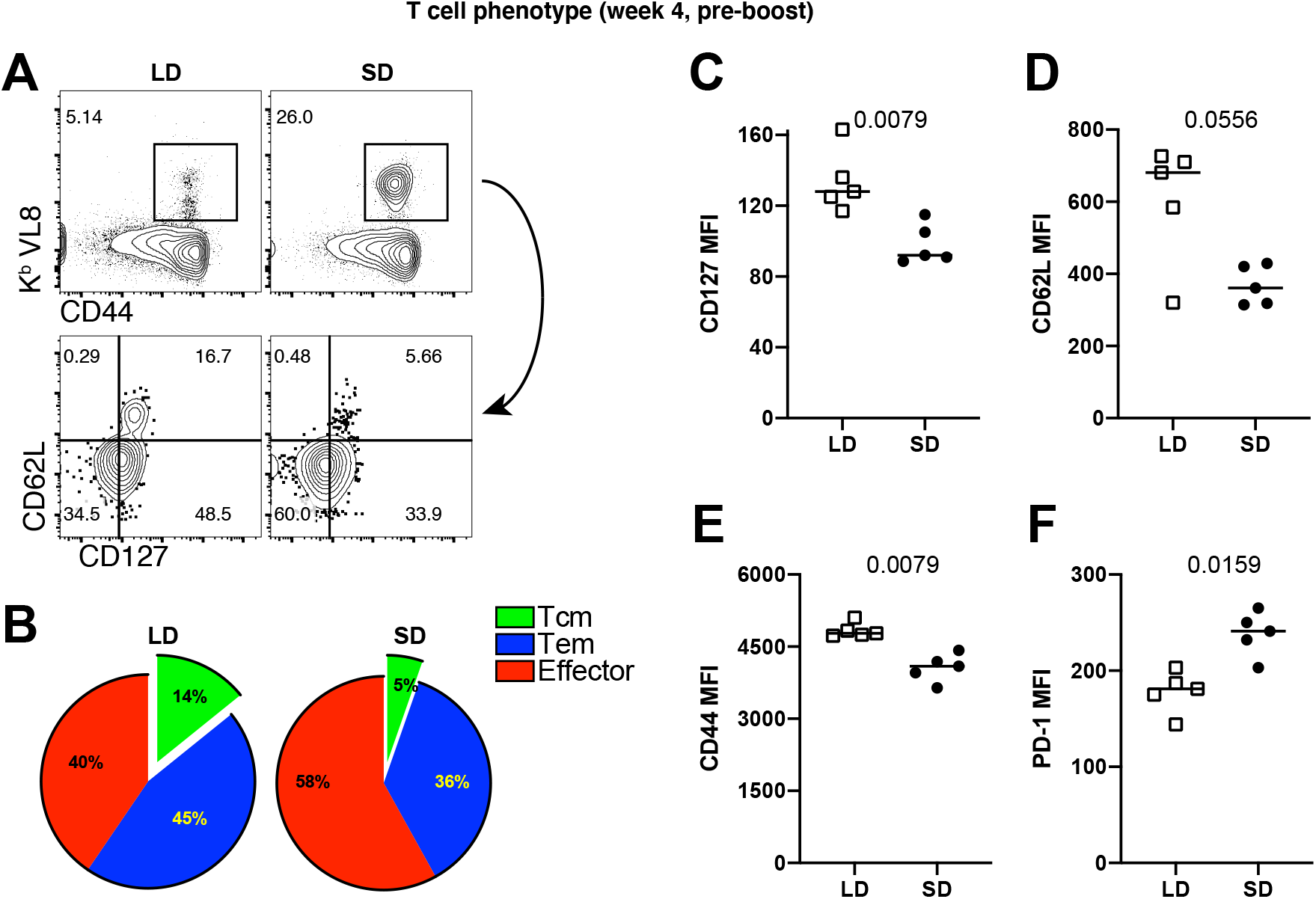
Phenotypic validation of CD8 T cell responses after a single prime with the Ad5-SARS-2 spike vaccine. **(A)** Representative FACS plots showing the frequencies of SARS CoV-2-specific CD8 T cells (K^b^ VL8) that differentiate into effector memory and central memory T cell subsets. **(B)** Summary of effector memory and central memory T cell subsets. **(C)** CD127 expression. **(D)** CD62L expression. **(E)** CD44 expression. **(F)** PD-1 expression. Panels B-F are gated from splenic SARS CoV-2-specific CD8 T cells (K^b^ VL8). All data are from day 28 post-prime. Data are from one experiment with n=5 per group. Experiment was repeated two additional times with similar results. Indicated *P* values were determined by Mann-Whitney U test.

Pre-existent immunity to adenoviral vectors can negatively affect vaccine-elicited immunity. We also reasoned that since mice that were primed with a SD harbor higher levels of virus-specific T cells and antibodies, this may result in more stringent competition for antigen during a subsequent booster immunization. To rule out these possibilities, we evaluated recall CD8 T cells in the absence of other pre-existing responses, by performing adoptive transfers of low numbers of purified SARS CoV-2-specific CD8 T cells into naïve hosts. Four weeks after priming mice with a LD or a SD, we FACS-purified splenic SARS CoV-2-specific CD8 T cells, and transferred these at equal low numbers (500 cells) into congenically distinct naïve mice. One day after adoptive transfer, all recipient mice were vaccinated with the SD (10^9^ PFU) of the SARS CoV-2 vaccine, and secondary CD8 T cell expansion was assessed by flow cytometry (Fig. 6A). We show that donor CD8 T cells from mice that were primed with a LD exhibited more robust secondary expansion compared to those primed with a SD (Fig. 6B-6E). These data show that a “gentle” antigenic prime elicits CD8 T cells that are intrinsically superior on a per-cell basis, and better able to mount a secondary response upon a booster immunization, irrespective of whether there are pre-existing vector or transgene-specific immune responses in the host. The adoptive transfer experiment above with normalized numbers of CD8 T cells was also consistent with our earlier scRNA-seq data showing that on a per-cell basis, a LD vaccine prime elicited less terminal differentiation with a superior Tcm differentiation.

**Figure 6.**
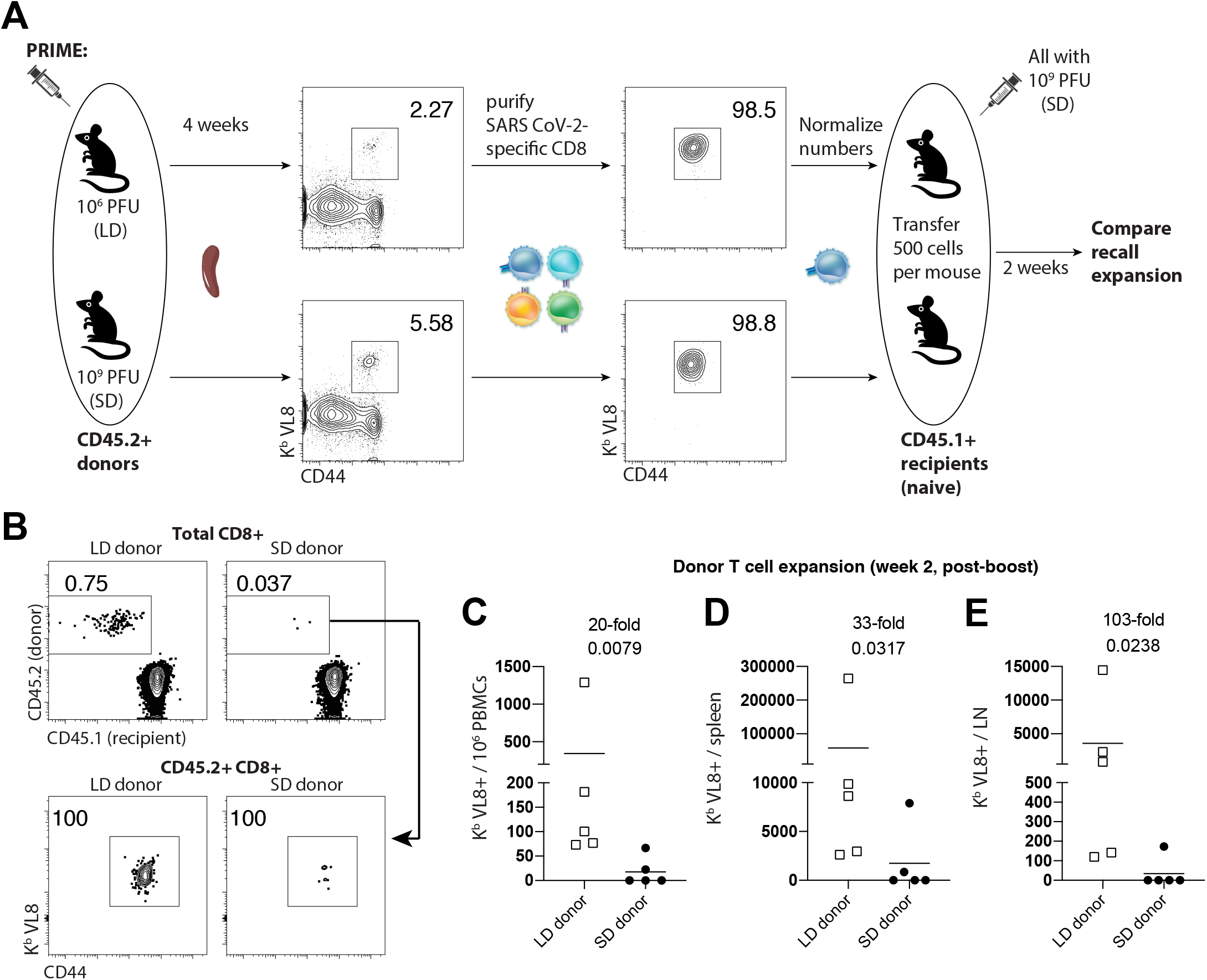
A LD prime elicits CD8 T cell responses with intrinsically superior anamnestic capacity. CD45.2+ mice were immunized intramuscularly with 10^6^ or 10^9^ PFU of Ad5-SARS-2 spike, and at day 28, splenic CD8 T cells were MACS-sorted. Subsequently, live, CD8+, CD44+, K^b^ VL8+ cells were FACS-sorted to ~99% purity, and numbers were normalized for adoptive transfer into CD45.1+ recipient mice. **(A)** Experimental approach for evaluating secondary expansion of donor CD8 T cells. **(B)** Representative FACS plots showing the frequencies of donor CD8 T cells after boosting. **(C)** Summary of donor-derived CD8 T cells in PBMCs. **(D)** Summary of donor-derived CD8 T cells in spleen. **(E)** Summary of donor-derived CD8 T cells in draining lymph nodes. Data from panels B-E are from day 14 post-boost. Data are from one experiment with n=5 per group. Experiment was repeated once with similar results. Indicated *P* values were determined by Mann-Whitney U test.

### LD/SD elicits superior antibody responses relative to SD/SD

Our analyses so far have been focused on T cell responses, but we also report profound differences in antibody responses. After a single prime immunization, the LD elicited antibody responses that were expectedly lower compared to the SD. However, antibody responses elicited by the LD expanded more robustly upon boosting (Fig. 7A). The LD/SD regimen exhibited superior germinal center B cell responses upon boosting, relative to the SD/SD regimen (Fig. 7B-7C). Generation of germinal center (GC) B cells after SARS CoV-2 vaccination is associated with potent SARS CoV-2 neutralizing antibody responses (*4*). Therefore, we performed neutralization assays using SARS CoV-2 pseudovirus to measure the ability of vaccine-elicited antibodies to block viral entry. Consistent with the profound increase in GC B cells, the LD/SD regimen elicited a 43-fold improvement in neutralizing antibodies compared to SD/SD (Fig. 7D).

**Figure 7.**
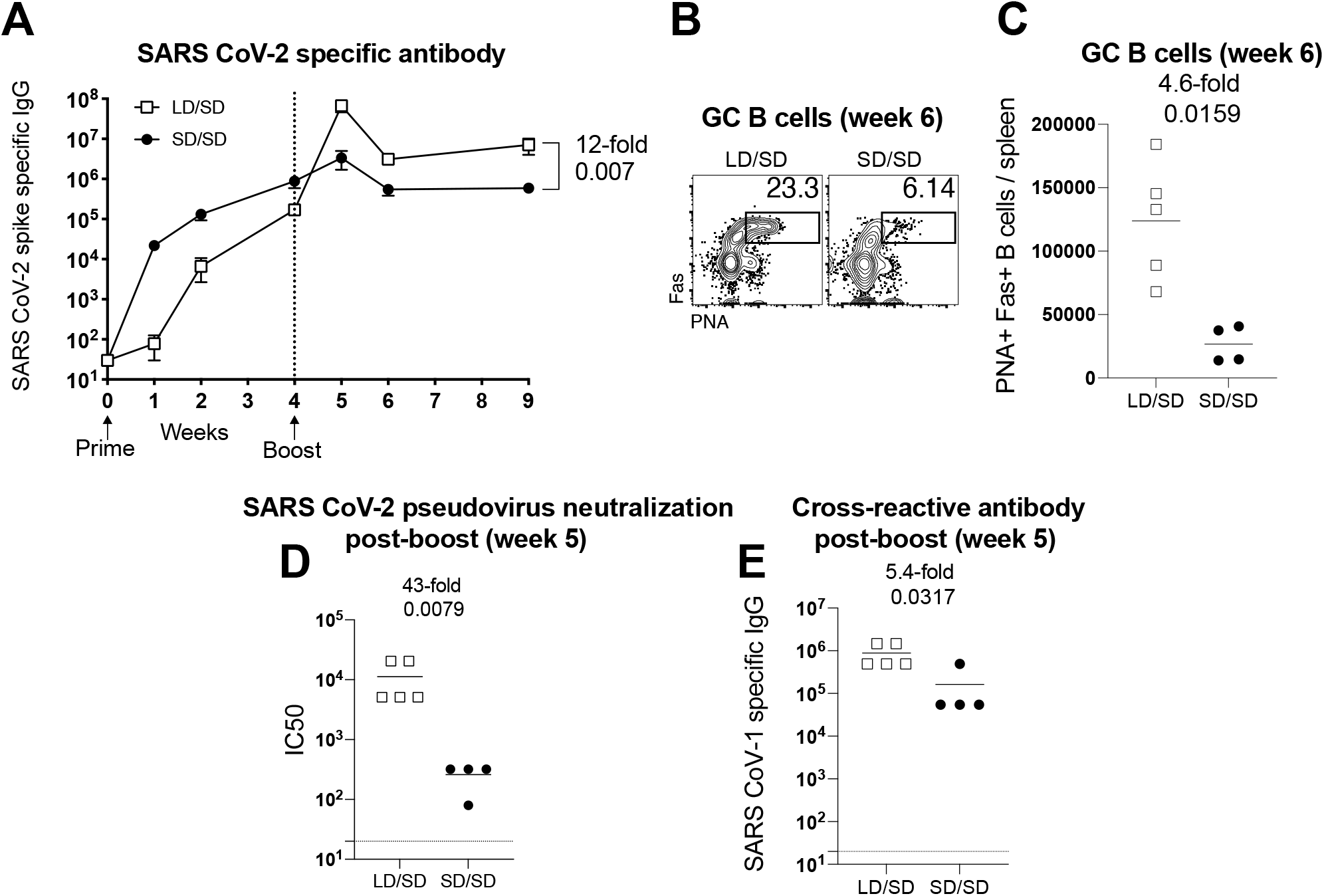
A LD/SD vaccine regimen elicits antibody responses with superior recall potential compared to a SD/SD vaccine regimen. Experimental approach was similar to Figure 1A. **(A)** Summary of SARS CoV-2 specific antibody responses in sera. **(B)** Representative FACS plots showing the frequencies of germinal center B cells in spleen (cells are gated on CD3-B220+ IgM-IgD-). **(C)** Summary of germinal center B cells in spleen. **(D)** Summary of SARS CoV-2 neutralizing antibody responses in sera. **(E)** Summary of cross-reactive (SARS CoV-1) antibody responses in sera. Data in panels D-E are from d7 post-boost (week 5). Data are from one experiment with n=4-5 per group. Experiment was repeated once with similar results. Indicated *P* values were determined by Mann-Whitney U test. Error bars represent SEM.

Currently, there are concerns about emerging SARS CoV-2 variants, and their potential to evade vaccine-elicited immune responses. A critical question is whether the current vaccines would protect against these variants or different coronaviruses that may appear in the future. We therefore interrogated whether changing the priming dose of our SARS CoV-2 vaccine could affect immune cross-reactivity against another coronavirus. To answer this simple question, we measured antibody responses to a distinct SARS coronavirus (original SARS CoV-1). Interestingly, the LD/SD regimen resulted in 5-fold improvement in cross-reactive antibody responses, relative to the SD/SD regimen (Fig. 7E). Altogether, these findings demonstrate that the anamnestic potential of antibody responses is also enhanced when the initial vaccine dose is reduced. In conclusion, we show that reducing the prime dose of an Ad5-based SARS CoV-2 vaccine confers an unprecedented advantage to T cell responses and antibody responses.

## Discussion

Adenovirus-based SARS CoV-2 vaccines are currently being deployed in humans to prevent COVID-19, including the Johnson & Johnson’s Janssen vaccine, AstraZeneca/Oxford vaccine, CanSino vaccine, and the Sputnik V vaccine. These vaccines have already been administered to millions of people, and have shown potent immunogenicity, safety, and efficacy against severe SARS CoV-2 infection. The CanSino and Sputnik V vaccines specifically utilize Ad5, which is the same vector platform used in our studies. Ad5 is among the most well-studied vaccine vectors, due in part to its extraordinary immunogenicity (*5, 6*). Although SARS CoV-2 vaccines prevent severe disease and death, they do not always confer sterilizing immunity, warranting further optimization of current vaccine regimens. Another issue is that there is still an insufficient number of vaccine doses available to immunize the entire population, motivating health authorities to administer half doses of vaccines in order to double the number of vaccinated people, but there is little knowledge of how vaccine fractionation could affect vaccine-elicited immunity.

Here, we show that fractionating the priming dose of an adenovirus-based SARS CoV-2 vaccine confers an unexpected immunological benefit. However, a possible trade-off of fractionating the priming dose is that it may initially result in lower immune responses, which may render the host transiently more susceptible to infection. Notwithstanding this potential concern, It is unclear what is the minimum level of immune responses required to protect against severe COVID-19, but a recent study demonstrated that even a low dose single-shot of an adenovirus-based SARS CoV-2 vaccine is sufficient to confer protection against severe disease in primates (*7*). Similarly, a single dose of a DNA-based SARS CoV-2 vaccine, which is ~1000-fold less immunogenic than a LD single-shot of our Ad5-based vaccine, was able to protect macaques from a SARS CoV-2 challenge (*8*). These prior data suggest low levels of immune responses are sufficient to protect against severe COVID-19, which can explain the extraordinary success of multiple experimental SARS CoV-2 vaccines in the last year.

There are historical examples of vaccine dose fractionation, also known as dose-sparing, as a way to allow more people to get vaccinated. For instance, prior studies have shown that administering a fifth of the recommended dose of the yellow fever virus vaccine results in comparable immune responses relative to the recommended dose (*9*). Similar results have been reported for influenza, measles, polio and typhoid vaccines (*10–12*), which has led to discussions of whether SARS CoV-2 vaccine doses should be fractionated to allow more people to get vaccinated. A rationale for fractionating SARS CoV-2 vaccines is that it may result in wider herd immunity even if vaccine efficacy is only partial, and that this could have a more striking effect in blocking transmission at the population level. However, an argument against vaccine fractionation is that it has not been rigorously tested with SARS CoV-2 vaccines.

Conceptually, our findings are contrary to conventional wisdom, because we show that when it comes to the immune system, sometimes “less is more.” Antigen is the first signal required for the activation of the adaptive immune system. However, the amount of antigen during a vaccine prime is not always proportional to the long-term magnitude and quality of the immune response, especially if the immune response is subsequently boosted. A prior study by Ahmed and colleagues in the lymphocytic choriomeningitis virus (LCMV) system showed that the duration of initial antigenic stimulation impacts T cell differentiation. This earlier study showed that CD8 T cells primed at the early stage of the infection preferentially differentiate into Tem, whereas CD8 T cells primed during the end of the infection preferentially differentiate into Tcm. Furthermore, we have previously shown that protracted acute infections favor Tem differentiation (*13*). Altogether, these prior reports suggest that a protracted antigen stimulation after a prime imprints a Tem phenotype, whereas a shortened antigen stimulation after a prime imprints a Tcm phenotype. Similar to this notion, we show that a LD prime favors Tcm differentiation, characterized by high expression of CD127, the IL-7 receptor a chain, which allows the T cell to survive long-term. Tcm CD8 T cells also exhibit enhanced recall capacity, which can explain the improved secondary expansion that was observed in our LD/SD regimen.

The LD/SD regimen also induced superior antibody responses compared to the SD/SD regimen, suggesting that a “gentle” antigen prime also generates memory B cell responses with significantly superior anamnestic capacity. Strikingly, the LD/SD regimen generated a 43-fold improved neutralizing response, compared to the SD/SD regimen. Thus, the effect of a LD prime on antibody responses is substantially more striking compared to its effect on T cell responses. Most of our mechanistic studies involve CD8 T cells, and our future studies will be aimed at understanding how a LD prime affects memory B cell differentiation. *In vitro* antibody neutralization is directly correlated with *in vivo* protection against SARS CoV-2 challenges (*8*), but we did not perform *in vivo* SARS CoV-2 challenges to compare immune protection, because in our pilot studies in k18-ACE2 mice, even a single-shot of our Ad5-SARS-2 spike vaccine provided near sterilizing protection; most animals that received a single shot of our Ad5-SARS-2 spike vaccine and were then challenged with SARS CoV-2 showed RNA levels that were undetectable or near the limit of detection. It is possible that our LD/SD regimen may confer a protective advantage relative to the SD/SD regimen at later times post-vaccination when vaccine-elicited responses wane. Therefore, in the long-term, a LD/SD regimen may potentially obviate the need for a third boost. The LD/SD regimen may also be useful to improve vaccine efficacy in individuals who develop suboptimal immune responses to vaccines, and who may benefit from having a higher level of immune responses.

There are growing concerns about new SARS CoV-2 variants and their ability to evade immune responses elicited by vaccination or natural infection. There are also concerns about re-emerging coronaviruses, such as SARS CoV-1, as well as novel coronaviruses that may enter the human population in the future, and an important question is whether the current vaccines would protect against future coronavirus pandemics. We investigated this issue of “immune coverage” by analyzing cross-reactive antibody responses (SARS CoV-1-specific), and we show that the LD/SD regimen resulted in improved cross-reactive antibody responses compared to SD/SD. Note also that the CD8 T cell response that we measured in these studies was specific for a highly conserved epitope (VNFNFNGL) that is present across multiple coronaviruses. There have been reports of re-infections caused by SARS CoV-2 variants, such as the B1.1.7 and B.1.351 variants, which are also thought to evade antibody responses elicited by vaccination (*14*). Interestingly, a study of recovered COVID-19 patients showed that virus-specific CD8 T cells are still able to recognize these variants, highlighting a critical role for CD8 T cells in providing broad protection against emerging variants (*15*). Altogether, the LD/SD regimen may be particularly effective in the context of universal coronavirus vaccines, because this regimen induces potent levels of cross-reactive immune responses.

The extent to which our results generalize to humans has not been determined, but a recent clinical trial with an adenovirus-based SARS CoV-2 vaccine (ChAdOx1, nCoV-19), in which the priming dose was accidentally reduced to half, reported superior efficacy relative to standard dose (90% efficacy for LD/SD, versus 62% efficacy for SD/SD) (*16*). However, in those studies it was unclear whether the improvement in the LD/SD group was due to prolonging the prime-boost interval or due to the priming dose itself. Our results bring more clarity to this confounding issue, as we show that the priming dose alone can have a substantial qualitative effect on immune responses. Historically, phase 1 vaccine trials compare a range of vaccine doses among different groups of individuals. However, they do not typically analyze the effect of dose-escalation within the same individual. The data presented here make a case for exploring intra-group vaccine dose escalation in future clinical trials. In summary, our results warrant a re-evaluation of current vaccine trial design, and most importantly, they may be useful to understand the effects of vaccine fractionation, which is a consideration during times of vaccine scarcity.

## ACKNOWLEDGEMENTS

We thank Drs. Susan Weiss, Stanley Perlman and Thomas Gallagher for discussions and reagents. This work was possible with a grant from the Emerging and Re-Emerging Pathogens Program (EREPP) at Northwestern University, and a grant from the National Institute on Drug Abuse (NIDA, DP2DA051912) to P.P.M.

## AUTHOR CONTRIBUTIONS

P.P.M., S.S., N.P., T.D. and T.C. designed and conducted the experiments. P.P.M. wrote the manuscript with feedback from all authors.

## Materials and Methods

### Mice and vaccinations

6-8-week-old C57BL/6 mice were used. Mice were purchased from Jackson laboratories (approximately half males and half females). Mice were immunized intramuscularly (50 μL per quadriceps) with an Ad5 vector expressing SARS CoV-2 spike protein (Ad5-SARS-2 spike) diluted in sterile PBS. Mice were housed at the Northwestern University Center for Comparative Medicine (CCM) in downtown Chicago. All mouse experiments were performed with approval of the Northwestern University Institutional Animal Care and Use Committee (IACUC).

### Reagents, flow cytometry and equipment

Single cell suspensions were obtained from PBMCs and various tissues as described previously (*17*). Dead cells were gated out using Live/Dead fixable dead cell stain (Invitrogen). SARS CoV-2 spike peptide pools used for intracellular cytokine staining (ICS) and these were obtained from BEI Resources. Biotinilated MHC class I monomers (K^b^ VL8, sequence VNFNFNGL) were obtained from the NIH tetramer facility at Emory University. The VNFNFNGL epitope is highly conserved among many multiple coronaviruses, representing a cross-reactive CD8 T cell response. It is located in position 539-546 of the SARS CoV-2 spike protein, or position 525-532 of the SARS CoV-1 spike protein. Cells were stained with fluorescently-labeled antibodies against CD8α (53-6.7 on PerCP-Cy5.5), CD44 (IM7 on Pacific Blue), TNFα (MP6-XT22 on PE-Cy7), IL-2 (JES6-5H4 on PE), IFNγ (XMG1.2 on APC), peanut agglutinin or PNA (conjugated to fluorescein), Fas (Jo2 on PE), IgD (11-26 on Pacific Blue), IgM (RMM-1 on PECy7), B220 (RA3-6B2 on PerCP-Cy5.5), and CD3 (145-2c11 on FITC). Fluorescently-labeled antibodies were purchased from BD Pharmingen, except for anti-CD44 (which was from Biolegend). Flow cytometry samples were acquired with a Becton Dickinson Canto II or an LSRII and analyzed using FlowJo (Treestar).

### Homologous (SARS CoV-2) and heterologous (SARS CoV-1) virus-specific ELISA

Binding antibody titers were measured using ELISA as described previously (*18, 19*), but using spike protein instead of viral lysates. In brief, 96-well flat bottom plates MaxiSorp (Thermo Scientific) were coated with 0.1μg/well of the respective spike protein, for 48 hr at 4°C. Plates were washed with PBS + 0.05% Tween-20. Blocking was performed for 4 hr at room temperature with 200 μL of PBS + 0.05% Tween-20 + bovine serum albumin. 6μL of sera were added to 144 μL of blocking solution in first column of plate, 1:3 serial dilutions were performed until row 12 for each sample and plates were incubated for 90 minutes at room temperature. Plates were washed three times followed by addition of goat anti-mouse IgG horseradish peroxidase conjugated (Southern Biotech) diluted in blocking solution (1:5000), at 100 μL/well and incubated for 90 minutes at room temperature. Plates were washed three times and 100 μL /well of Sure Blue substrate (Sera Care) was added for approximately 8 minutes. Reaction was stopped using 100 μL/well of KPL TMB stop solution (Sera Care). Absorbance was measured at 450 nm using a Spectramax Plus 384 (Molecular Devices). SARS CoV-2 spike protein was made at the Northwestern Recombinant Protein Production Core by Dr. Sergii Pshenychnyi using a plasmid that was produced under HHSN272201400008C and obtained through BEI Resources, NIAID, NIH: Vector pCAGGS Containing the SARS-Related Coronavirus 2, Wuhan-Hu-1 Spike Glycoprotein Gene (soluble, stabilized), NR-52394. SARS CoV-1 spike protein was obtained through BEI Resources, NIAID, NIH: SARS-CoV Spike (S) Protein deltaTM, Recombinant from Baculovirus, NR-722.

### Pseudovirus neutralization assays

A SARS CoV-2 Spike pseudotyped lentivirus kit was obtained through BEI Resources, NIAID, NIH (SARS-Related Coronavirus 2, Wuhan-Hu-1 Spike-Pseudotyped Lentiviral Kit V2, NR-53816). We used Human Embryonic Kidney Cells (HEK-293T) expressing Human Angiotensin-Converting Enzyme 2, also known as HEK-293T-hACE2, which are susceptible to SARS CoV-2. This cell line was obtained through BEI Resources, NIAID, NIH NR-52511. Serial dilutions of sera were incubated with the SARS CoV-2 Spike pseudotyped lentivirus, following a protocol by Balazs and Bloom (*20*). Cells were lysed using luciferase cell culture lysis buffer (Promega). Luciferase reaction was performed using 30 μL of cell lysis (Promega). The reaction was added to 96-well black optiplates (Perkin Elmer). Luminescence was measured using Perkin Elmer Victor^3^ luminometer.

### Ad5-SARS-2 spike vaccine

This non-replicating Ad5 vector is E1/E3 deleted and expresses the SARS CoV-2 spike protein (strain 2019-nCoV-WIV04) within the putative E1 site. This vector contains a CMV (Cytomegalovirus) promoter driving the expression of SARS-CoV-2-Spike protein. These vaccines were propagated on trans-complementing HEK293 cells (ATCC), purified by cesium chloride density gradient centrifugation, titrated, and then frozen at −80 °C.

### Single cell RNA-Seq Data Acquisition and Analysis

C57BL/6 mice were immunized with a LD (10^6^ PFU) or a SD (10^9^ PFU) of Ad5-SARS-2 spike, and at day 28, splenic CD8 T cells were MACS-sorted with a MACS negative selection kit (STEMCELL). Purified CD8 T cells were stained with K^b^ VL8 tetramer, live dead stain, and flow cytometry antibodies for CD8 and CD44 to gate on activated CD8 T cells. Live, CD8+, CD44+, K^b^ VL8+ cells were FACS-sorted to ~99% purity on a FACS Aria cytometer (BD Biosciences) and delivered to the Northwestern University NUSeq core for scRNA-seq using Chromium NextGem 5’ v2 kit (10X Genomics). After the library was sequenced, the output file in BCL format was converted to fastq files and aligned to mouse genome in order to generate a matrix file using the Cell Ranger pipeline (10X Genomics). These upstream QC steps were performed by Drs. Ching Man Wai and Matthew Schipma at the Northwestern University NUSeq core. Further analyses were performed in R using the Seurat package v4.0 (PMID: 30638736), as previously described (*21*). Terminal effector gene signatures were derived using the edgeR package (*22*), comparing effector memory to terminal effector CD8 T cells (*3*). Clusters representing less than 4% of each population were excluded from downstream analyses. TCR analyses was performed using the scRepertoire package (*23*). Only cells expressing both TCRa and TCRb chains were selected. For cells with more than 2 TCR chains, only the top 2 expressed chains were used.

### Statistical analysis

Statistical analyses used the Mann Whitney test. Dashed lines in data figures represent limit of detection. Data were analyzed using Prism (Graphpad).

